# Evaluation of a potential redox switch in blood coagulation tenase

**DOI:** 10.1101/515718

**Authors:** Kristina M. Cook, Diego Butera, Philip J. Hogg

## Abstract

Blood coagulation factor IXa (FIXa) activates factor X that leads to thrombin and fibrin formation and a stable thrombus. FIXa catalytic efficiency is markedly enhanced when bound to the co-factor, factor VIII (FVIII), and a negatively charged phospholipid surface in the tenase complex. Small redox active peptides and protein oxidoreductases have been shown previously to have some FIXa co-factor activity and thiol modifying agents have been reported to influence FVIII activity. These observations suggested that FIXa might contain an allosteric disulfide that is regulated by FVIII. This idea was tested by measuring the influence of FVIII on the redox state of FIXa disulfide bonds and the effect of plasma oxidoreductases on FIXa activity. The redox state of nine of the 11 disulfide bonds in FIXa was measured using differential cysteine labelling and mass spectrometry and all were oxidized in the protein, and this did not change upon binding of the enzyme to FVIII. All eight disulfide bonds in FVIII were also predominantly oxidized and this did not appreciably change upon FIXa binding. In addition, relevant protein reductants in the circulation inhibited rather than activated FIXa activity. In conclusion, we found no evidence that the co-factor function of FVIII involves a change in the redox state of one or more FIXa disulfide bonds.

## Introduction

Cleavage of peptide bonds controls a number of key processes in the mammalian circulation, such as blood coagulation and complement activation. Disulfide bonds are the second most frequent covalent bond, after the peptide bond, linking the polypeptide backbone of proteins, and their cleavage is emerging as another important mechanism of regulation of blood proteins (Butera et al., 2014).

It has been estimated that the 2,000 or so human plasma proteins contain about 10,000 disulfide bonds (Butera et al., 2014). Most of these disulfide bonds remain unchanged for the life of the protein, but some, the allosteric disulfides, control the function of the mature protein in which they reside when they are cleaved (Cook and Hogg, 2013). About 30 proteins have been shown to be regulated by allosteric disulfide bonds to date, and 12 of these are found in the mammalian circulation (Butera et al., 2014; Cook and Hogg, 2013; Hogg, 2013). One of these is hTryptase β, a serine protease secreted by mast cells.

Tryptases are by far the most abundant constituents of the mast cells granules, and these serine proteases are involved in many acute and chronic inflammatory processes. The Cys220-Cys248 disulfide bond in hTryptase-β and its ortholog, mouse mast cell protease-6, exists in oxidized and reduced states in the enzyme stored and secreted by mast cells (Cook et al., 2013). The different redox states of hTryptase-β have different specificity and catalytic efficiency for hydrolysis of protein substrates. The tryptases belong to the serine protease subfamily S1A, a large group of trypsin-like, neutral serine proteases involved in coagulation, immunity and digestion (Yousef et al., 2004). The Cys191-Cys220 disulfide bond (chymotrypsin numbering) is conserved in all S1A serine proteases, except for the cathepsins and granzymes (Yousef et al., 2004).

The structural/functional role of the Cys191-Cys220 disulfide bond has been investigated in some blood coagulation proteases. The disulfide bond in FXIIa (Cys559-Cys590 in FXII) is essential for activity, as mutation of Cys571 to serine in the Hageman trait blood coagulation disorder inactivates the enzyme (Miyata et al., 1989). Similarly, ablation of the disulfide blood in FVIIa (Cys400-Cys428 in FVII) (Higashi et al., 1997) and thrombin (Cys564-Cys594 in prothrombin) (Bush-Pelc et al., 2007) markedly impairs activity. We have explored herein a possible allosteric role for the equivalent disulfide bond in blood coagulation FIXa (Cys407-Cys435 in FIX) (Fig. 1).

**Figure 1.**
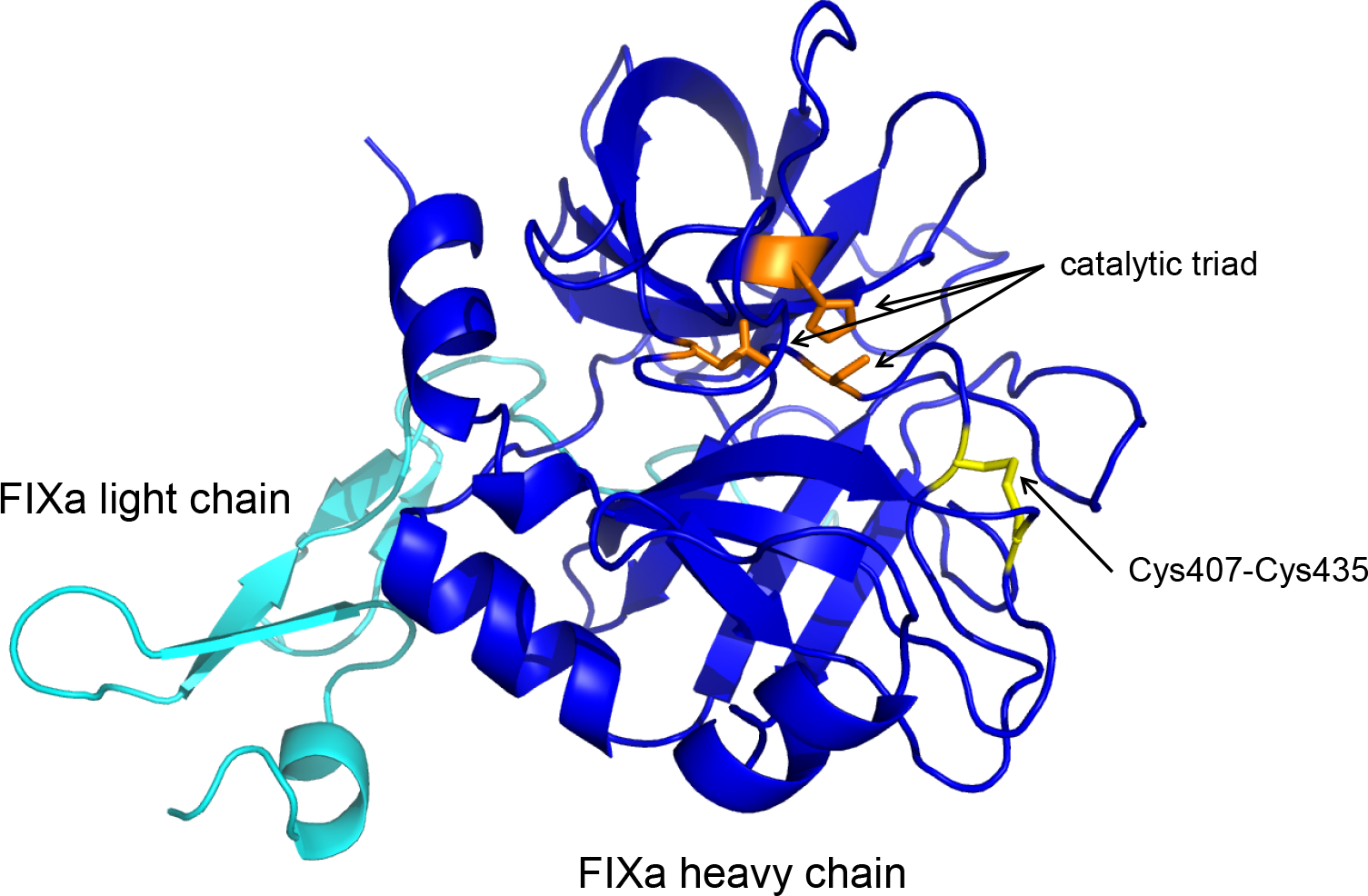
Ribbon structure of human FIXa showing the heavy and light chains in blue and cyan, respectively, the active site catalytic triad in orange, and the Cys407-Cys435 disulfide bond in yellow (PDB identifier 1rfn) (Hopfner et al., 1999).

FIXa activates FX in a complex with cofactor FVIII and phospholipid called tenase. FX activation leads to thrombin and fibrin formation, which stabilize the developing thrombus. In a study designed to identify cofactor requirements for FVIII, a tetrapeptide with the same sequence as the active-site of the protein reductant thioredoxin, CGPC, was found to activate FX in a FIXa-dependent manner (Bayele et al., 2010). The oxidoreductases, thioredoxin and protein disulfide isomerase (PDI), were also found to possess some co-factor activity, although were much less efficient than FVIII. These observations suggested that FIXa might contain an allosteric disulfide, possibly the Cys407-Cys435 bond, which is regulated by FVIII.

In this study, we have characterized the influence of FVIII on the redox state of FIXa disulfide bonds and the effect of oxidoreductases on FIXa activity. We found that the redox state of none of the FIXa and FVIII disulfide bonds was influenced by interaction of the proteins, and plasma protein reductants inhibited rather than activated FIXa activity. We conclude that the co-factor function of FVIII does not involve a change in the redox state of one or more FIXa disulfide bonds, at least in our system.

## Results

### Redox state of the FIXa disulfide bonds before and after incubation with FVIII

Purified human plasma FIXa and recombinant human FVIII were employed in our reactions. The co-factor function of FVIII was tested by incubating phospholipid vesicles with FIXa and FX in the absence or presence of increasing concentrations of FVIII, and measuring formation of FXa from hydrolysis of a peptide p-nitroanilide substrate (Fig. 2). Having confirmed the activity of our FIXa and FVIII, the redox state of the FIXa and FVIII disulfide bonds were then measured using differential cysteine labelling and mass spectrometry.

**Figure 2.**
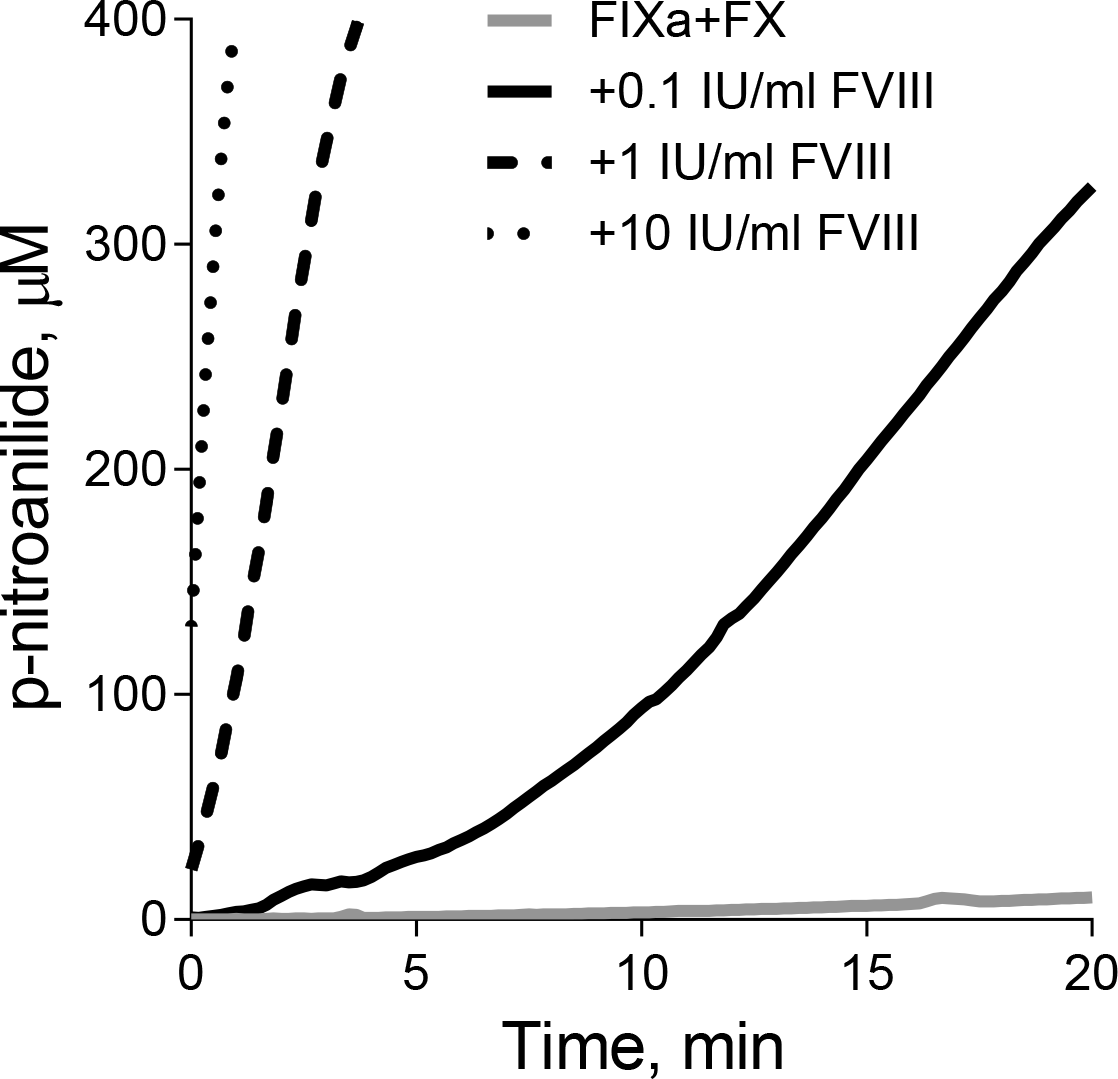
Co-factor activity of FVIII in our system. Phospholipid vesicles (58 μM) were incubated with FIXa (1 nM) and FX (240 nM) in the absence or presence of FVIII (0.1, 1 or 10 IU/ml) for 2 min, and formation of FXa was monitored by hydrolysis of the chromogenic substrate, Suc-Ile-Glu(γPip)Gly-Arg-pNA (500 μM). The solid lines are the average of 3 experiments. There was negligible hydrolysis of the chromogenic substrate in the absence of FX.

Differential cysteine labeling and mass spectrometry allows for an estimation of the fraction of a disulfide bond that is reduced in the protein preparation. Reduced disulfide bond cysteines are alkylated with 2-iodo-N-phenylacetamide (^12^C-IPA) and the oxidized disulfide bond cysteines with a stable carbon-13 isotope of IPA (^13^C-IPA) following reduction with dithiothreitol (Fig. 3A). The ratio of labeling of peptides containing the disulfide bond cysteines with ^12^C-IPA compared to ^13^C-IPA represents the fraction of the disulfide in the population that is in the reduced state. For example, a tandem mass spectrum of the FIXa peptide, HEGGRDS<u>C</u>QGDSGGPH, containing ^13^C-IPA-labeled Cys407 is shown in Fig. 3B. The advantage of this pair of cysteine alkylators is that they have the same chemical reactivity and the same structure, which enhances the reliability of alkylation and resolution, and detection of the alkylated peptides by mass spectrometry (Pasquarello et al., 2004). A mass difference of 6 Da is the only change in a peptide labeled with ^12^C-IPA or ^13^C-IPA.

**Figure 3.**
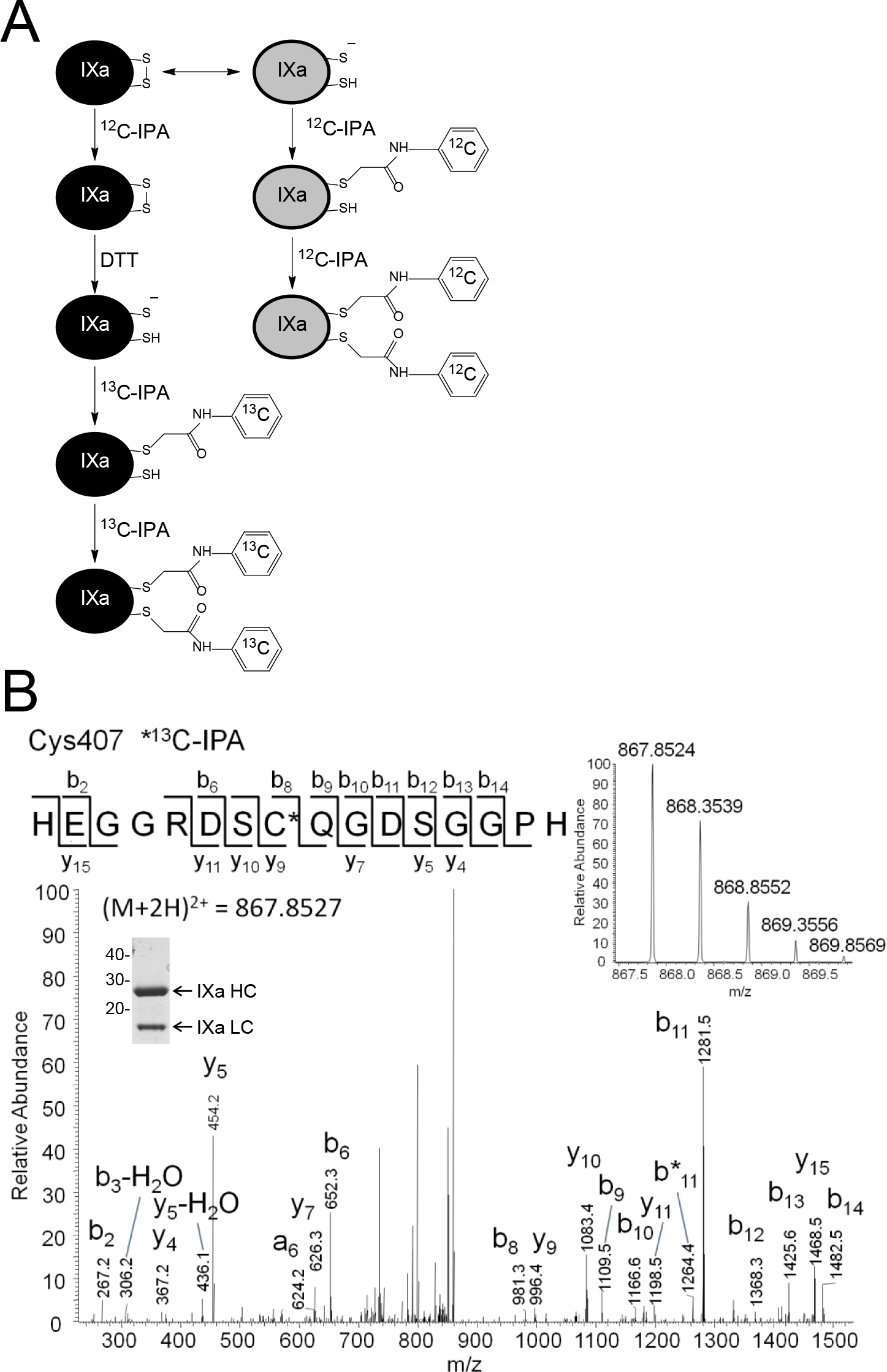
Differential cysteine labeling and mass spectrometry of FIXa. **A**. Reduced disulfide bond cysteines in FIXa were alkylated with ^12^C-IPA and the oxidized cysteine thiols with ^13^C-IPA following reduction with DTT. One or both of the reduced disulfide bond cysteines can be labeled with either alkylator. The ratio of ^12^C-IPA to ^13^C-IPA labeling represents the proportion of the disulfide bonds in the population that are in the reduced state. **B**. Tandem mass spectrum of the HEGGRDS<u>C</u>QGDSGGPH peptide showing Cys407 (underlined) labeled with ^13^C-IPA (disulfide bonded cysteine). The accurate mass spectrum of the peptide is shown in the inset (observed [M+2H]^2+^ = 867.8527 m/z; expected [M+2H]^2+^ = 867.8527 m/z). Also shown as an inset is the FIXa resolved on reducing SDS-PAGE and stained with colloidal coomassie. The positions of molecular weight standards in kDa are shown at the left.

FIXa contains 11 disulfide bonds comprising three in the heavy chain, seven in the light chain and one interchain bond. The redox state of nine of the 11 FIXa disulfide bonds was determined (Table 1). Peptides containing the light chain cysteines, Cys97, Cys108, Cys102 or Cys117, were not resolved in the analysis so their redox state could not be defined. Up to 15 tryptic or chymotryptic peptides containing one or both of the other 18 cysteines involved in nine disulfide bonds were resolved, which allowed for more than one estimate of the redox state in most cases (Table 1).

**Table 1.** Redox state of the FIXa disulfide bonds before and after incubation with equimolar FVIII.

| FIXa disulfide bonds | FIXa |  |  | FIXa+FVIII |  |  |
| --- | --- | --- | --- | --- | --- | --- |
|  | % oxidized<sup>a</sup> | % reduced<sup>a</sup> | number of peptides | % oxidized | % reduced | number of peptides |
| Cys64-Cys69 | 86 | 14 | 1 | 0 | 100 | 1 |
| Cys97-Cys108 | n.d.<sup>b</sup> | n.d. | 0 | n.d. | n.d. | 0 |
| Cys102-Cys117 | n.d. | n.d. | 0 | n.d. | n.d. | 0 |
| Cys119-Cys128 | 100 | 0 | 2 | 99 | 1 | 2 |
| Cys134-Cys145<sup>c</sup> | 98-100 | 0-2 | 6 | 98-100 | 0-2 | 10 |
| Cys141-Cys155 | 100 | 0 | 2 | 99 | 1 | 3 |
| Cys157-Cys170 | 100 | 1 | 2 | 99 | 1 | 2 |
| Cys178-Cys335 | 98 | 2 | 12 | 99 | 1 | 17 |
| Cys252-Cys268 | 100 | 0 | 15 | 100 | 0 | 17 |
| Cys382-Cys396 | 99 | 1 | 14 | 100 | 0 | 17 |
| Cys407-Cys435 | 97 | 3 | 14 | 97 | 3 | 15 |

a Average percent of alkylation of tryptic and/or chymotryptic peptides analysed containing either one or both of the disulfide bond cysteines.
b No peptides containing this Cys were resolved in the analysis.
c Peptides containing Cys134 and/or Cys145 only were not resolved in the analysis. These Cys were resolved on peptides containing Cys from other disulfide bonds, namely Cys128 and Cys141. A range of oxidized and reduced Cys134-Cys145 disulfide bond is presented based on the redox status of the Cys119-Cys128 and Cys141-Cys155 disulfides that were determined separately.

With one exception, all nine disulfide bonds were oxidized in the human plasma protein. Only a few percent of the bonds were found to be reduced, which is at the limit of detection for this technique. Cys69 of the light chain Cys64-Cys69 disulfide bond was resolved on 1 tryptic peptide (residues 69-75). Fourteen percent of this disulfide bond was calculated to be reduced, although this number should be treated with caution as no alternative peptides were resolved to validate this estimate.

FIXa was then incubated with an equimolar concentration of FVIII for 1 h, unpaired cysteines in the reaction were frozen by alkylation with ^12^C-IPA, and the redox state of FIXa disulfide bonds re-analyzed. There was no appreciable change in the redox state of the nine FIXa disulfide bonds upon interaction with FVIII (Table 1).

### Redox state of the FVIII disulfide bonds before and after incubation with FIXa

The redox state of the eight FVIII disulfide bonds (McMullen et al., 1995) was also determined in the analysis. All eight bonds were predominantly oxidized in the recombinant protein (Table 2). Less than 10% of any of the disulfides were found to be reduced in the protein preparation. Incubation of the FVIII with equimolar FIXa for 1 h did not appreciably change the redox state of the eight disulfide bonds (Table 2). Five of the eight disulfide bonds appeared to be more reduced following incubation with FIXa, although not obviously so.

**Table 2.**
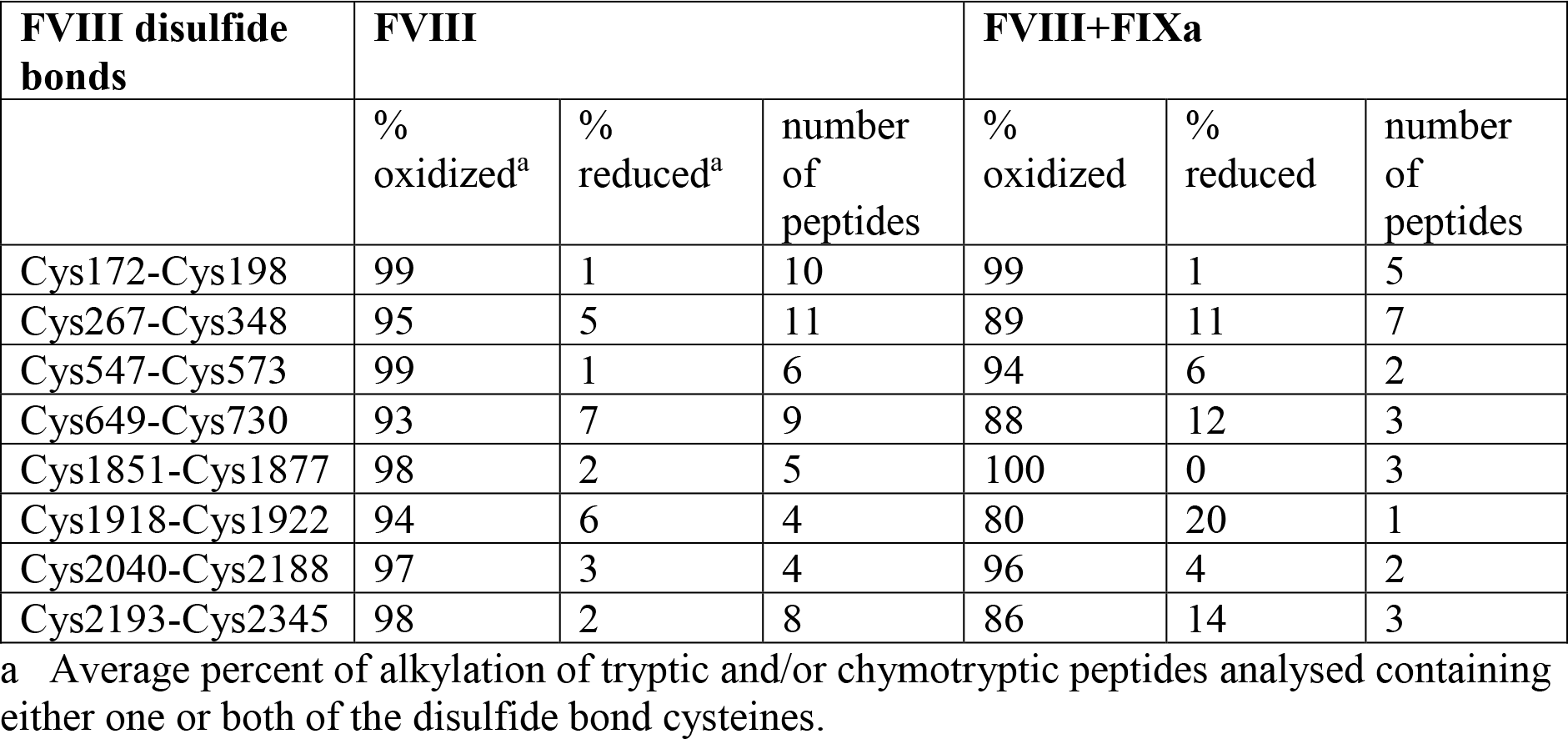
Redox state of the FVIII disulfide bonds before and after incubation with equimolar FIXa.

### Evaluation of the co-factor activity of protein reductants in activation of FX by FIXa

The co-factor activity of thioredoxin and PDI in FIXa activation of FX was determined in two assay formats: the COAMATIC FVIII assay to replicate the conditions employed by Bayele et al. (Bayele et al., 2010), and an assay where the individual tenase reactants were reconstituted.

Low micromolar concentrations of thioredoxin or PDI, which exceed the levels of these proteins in plasma, inhibited FIXa activation of FX in the COAMATIC assay by ~70% (Fig. 4A). The same result was observed when the individual components of the tenase complex were reconstituted (Fig. 4B). To test whether the reductants were inhibiting FIXa in the system, rather than FVIII or FX, the FIXa was incubated with thioredoxin or PDI first and the reductants quenched by alkylating the active site dithiols with N-ethylmaleimide. The treated FIXa was added to the other tenase components and formation of FXa measured. The same level of inhibition of FX activation was observed as when the reductants were added with all tenase components (Fig. 4C). This result indicates that thioredoxin or PDI are inhibiting FIXa in the assay.

**Figure 4.**
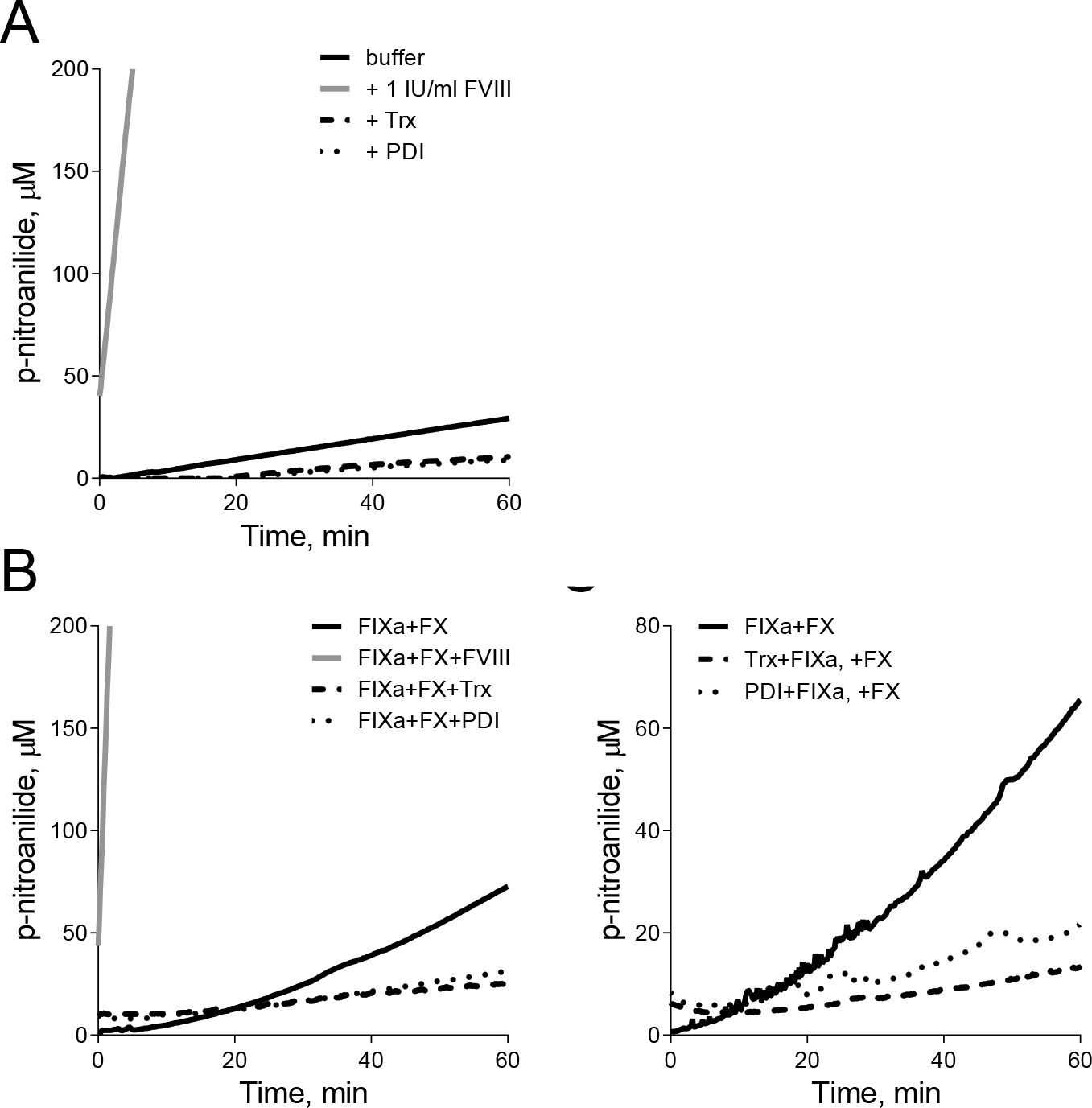
Evaluation of the co-factor activity of protein reductants in activation of FX by FIXa. A. Co-factor activity of thioredoxin (Trx) and PDI in the COAMATIC assay. Assay components, according to the manufacturers protocol, were incubated with either buffer, plasma, FVIII (1 IU/ml), Trx (24 μM) or PDI (31 μM) for 2 min, and formation of FXa was monitored by hydrolysis of the chromogenic substrate. The solid lines are the average of 3 experiments. B. Co-factor activity of Trx and PDI in an assay using purified components. Phospholipid vesicles (58 μM) were incubated with FIXa (1 nM) and FX (240 nM) in the absence or presence of FVIII (1 IU/ml), Trx (16 μM), or PDI (28 μM) for 2 min, and formation of FXa was monitored by hydrolysis of the chromogenic substrate, Suc-Ile-Glu(γPip)Gly-Arg-pNA (500 μM). The solid lines are the average of 3 experiments. There was negligible hydrolysis of the chromogenic substrate in the absence of FX. C. FIXa (1 nM) was incubated with Trx (16 μM) or PDI (28 μM) for 10 min and unpaired cysteine thiols in the reactions were alkylated with N-ethylmaleimide (10 mM) for 10 min. Phospholipid vesicles (58 μM) and FX (240 nM) were added and incubated for 2 min, then formation of FXa was monitored by hydrolysis of the chromogenic substrate. The solid lines are the average of 3 experiments.

## Discussion

Changing the redox environment of plasma by adding reduced or oxidized glutathione alters clotting times (Bayele et al., 2002). The mechanism(s) of this effect are not known and the physiological relevance is uncertain. It has been proposed that the co-factor function of FVIII in the tenase complex may involve thiol/disulfide exchange in FIXa (Bayele et al., 2010), and this is the hypothesis that we tested.

We investigated potential thiol/disulfide exchange in FIXa and/or FVIII by measuring the redox state of the disulfide bonds in both proteins before and after interaction of the factors. Nine of the 11 disulfide bonds in FIXa and all eight disulfide bonds in FVIII were resolved in the analysis and none were found to be significantly reduced before or after interaction. Two FIXa disulfide bonds in the light chain were not identified in the analysis and it is possible that they are redox active. We can conclude that thiol/disulfide exchange does not appear to be required for efficient tenase activity, at least in our *in vitro* setting using purified proteins. We cannot rule out thiol/disulfide exchange events in tenase function *in vivo*.

Thiol modifying agents have been reported to affect FVIII activity *in vitro* and *in vivo*. FVIII activity is variably affected by treatment with thiol alkylating or oxidizing agents *in vitro* (Austen, 1970; Kaelin, 1975; Knudsen et al., 2005; Manning et al., 1995), while infusion of the small thiol, N-acetylcysteine, into healthy subjects increased plasma FVIII activity (Knudsen et al., 2005). These findings imply a role for unpaired cysteine thiols in FVIII activity. In contrast to these reports, we found no effect of micromolar concentrations (500 μM) of the thiol alkylators, N-ethyl maleimide, iodoacetamide and methyl methanethiosulfonate, on FVIII activity in our tenase assay (data not shown).

We tested the reported co-factor activity of thioredoxin and PDI in the tenase assay format used by Bayele et al. (Bayele et al., 2010), and an assay where the individual tenase reactants were reconstituted. Both oxidoreductases inhibited rather than promoted FIXa activity in both assays. Our result is consistent with an early study showing that micromolar concentrations of thioredoxin rapidly inactivate FIX and FX (Savidge et al., 1979).

It is possible that thioredoxin and PDI are reducing the Cys407-Cys435 disulfide in factor FIXa, which is the same conserved bond that is required for efficient FXIIa (Miyata et al., 1989), FVIIa (Higashi et al., 1997) and thrombin (Bush-Pelc et al., 2007) activity. Fifteen disulfide bond oxidoreductases have been identified in plasma and/or on the surface of platelets or leukocytes (Butera et al., 2014; Lackman et al., 2007), so there is capacity for reduction of FIXa in the circulation. There is currently no *in vivo* evidence for regulation of FIXa activity by one or more of these oxidoreductases, although this merits further study.

## Materials and Methods

### Redox state of the FIXa and FVIII disulfide bonds

The redox state of the human plasma FIXa (Enzyme Research Laboratories) (Fig. 2A) and recombinant human FVIII (Advate, Baxter Healthcare) disulfide bonds were determined by differential cysteine labeling and mass spectrometry. The analysis of was performed as we have described (Cook et al., 2013), with the following modifications. Unpaired cysteine thiols in FIXa, FVIII, and an equimolar mixture of the two proteins incubated for 1 h at room temperature in phosphate-buffered saline, were alkylated with 10 mM 2-iodo-N-phenylacetamide (^12^C-IPA, Cambridge Isotopes) for 1 h at room temperature. The proteins were resolved on SDS-PAGE and stained with SYPRO Ruby (Invitrogen). Thiols were labelled as described (Butera et al., 2018; Read et al., 2017). Briefly, the FIXa or FVIII bands were excised, destained, dried, and incubated with 100 mM dithiothreitol (DTT), before incubation with 10 mM 2-iodo-N-phenylacetamide where all 6 carbon atoms of the phenyl ring have a mass of 13 (^13^C-IPA) (Cambridge Isotopes). The slices were washed and dried before digestion of proteins with 20 ng/μl of trypsin (Promega) or 12 ng/μl of chymotrypsin (Roche) in 25 mM NH_4_CO_2_ overnight at 25 °C. Peptides were eluted from the slices with 5% formic acid, 50% acetonitrile. Liquid chromatography, mass spectrometry and data analysis were performed as described (Cook et al., 2013).

The redox state of the FIXa and FVIII disulfide bonds were estimated from the relative ion abundance of peptides containing disulfide cysteines alkylated with either ^12^C-IPA or ^13^C-IPA. Relative ion abundance of peptides was calculated using the area of the peak detected from the extracted ion chromatograms generated by the XCalibur Qual Browser software (Version 2.0.7, Thermo Scientific). The ratio of ^12^C-IPA to ^13^C-IPA labeling represents the fraction of the cysteine in the population that is in the reduced state.

### FX activation assays

Recombinant human thioredoxin-1 was produced as previously described (Schwertassek et al., 2007a; Schwertassek et al., 2007b). The cDNA encoding full-length human PDI (residues 1-508) was cloned into the *Bam*H *I* and *Eco*R *I* sites of pAE vector (Ramos et al., 2004) to generate an N-terminally 6xHis-tagged PDI protein. The integrity of the construct was confirmed by automated sequencing. Expression and purification of PDI was performed by the Protein Production Unit at Monash University, Clayton, Victoria. Briefly, the pAE-PDI construct was transfected into *E.coli* BL21 (DE3)C41 cells and protein expressed in autoinduction media at 30°C for 4 h. The cells were lysed by sonication and PDI in the soluble fraction was purified by nickel affinity chromatography followed by gel filtration on Sephacryl S-75. Protein purity of 95-97% was achieved. The active site disulfide bonds of the reductants were reduced prior to use by incubating with 10 mM DTT for 30 min and removal of the unreacted DTT with a Zeba spin desalting column (Thermo Scientific). 1,2-Dioleoyl-*sn*-glycero-3-phosphocholine, 1,2-dioleoyl-*sn-*glycero-3-phospho-L-serine and 1,2-dioleoyl-*sn*-glycero-3-phosphoethanolamine in chloroform (Avanti Polar Lipids Inc) were combined in the molar ratio 60:20:20, respectively, and the solvent evaporated under nitrogen gas. The lipids were dissolved in Hepes buffered saline and unilamellar vesicles prepared by sonication.

The co-factor activity of thioredoxin and PDI in FIXa activation of FX was determined in two assay formats. The COAMATIC FVIII assay (Chromogenix) was performed as described by the manufacturer. In the other assay, the individual reactants were reconstituted. Phospholipid vesicles (58 μM) were incubated with FIXa (1 nM) and human plasma FX (240 nM, Enzyme Research Laboratories) in the absence or presence of FVIII (1 IU/ml), thioredoxin (16 μM), or PDI (28 μM) for 2 min in 50 mM Hepes, pH 7.4 buffer containing 100 mM NaCl, 5 mM CaCl_2_, 0.1 mM MnCl_2_ and 0.5% bovine serum albumin. The chromogenic substrate, Suc-Ile-Glu(γPip)Gly-Arg-pNA (500 μM, Hyphen Biomed), was added and FXa formation monitored from the release of p-nitroanilide measured by absorbance at 405 nm using a kinetic microplate reader.

## Acknowledgments

Funding was provided by the National Health and Medical Research Council of Australia (PJH).

## List of Abbreviations

FIX: factor IX
FIXa: activated factor IX
FVIII: factor VIII
FX: factor X
FXa: activated factor X
Trx: thioredoxin

## References

Austen, D.E. (1970). Thiol groups in the blood clotting action of factor VIII. Br J Haematol 19, 477–484.

Bayele, H.K., Murdock, P.J., and Pasi, K.J. (2010). Residual factor VIII-like cofactor activity of thioredoxin and related oxidoreductases. Biochim Biophys Acta 1800, 398–404.

Bayele, H.K., Murdock, P.J., Perry, D.J., and Pasi, K.J. (2002). Simple shifts in redox/thiol balance that perturb blood coagulation. FEBS Lett 510, 67–70.

Bush-Pelc, L.A., Marino, F., Chen, Z., Pineda, A.O., Mathews, F.S., and Di Cera, E. (2007). Important role of the cys-191 cys-220 disulfide bond in thrombin function and allostery. J Biol Chem 282, 27165–27170.

Butera, D., Cook, K.M., Chiu, J., Wong, J.W., and Hogg, P.J. (2014). Control of blood proteins by functional disulfide bonds. Blood 123, 2000–2007.

Butera, D., Passam, F., Ju, L., Cook, K.M., Woon, H., Aponte-Santamaria, C., Gardiner, E., Davis, A.K., Murphy, D.A., Bronowska, A., et al. (2018). Autoregulation of von Willebrand factor function by a disulfide bond switch. Sci Adv 4, eaaq1477.

Cook, K.M., and Hogg, P.J. (2013). Post-translational control of protein function by disulfide bond cleavage. Antioxid Redox Signal 18, 1987–2015.

Cook, K.M., McNeil, H.P., and Hogg, P.J. (2013). Allosteric control of bII-tryptase by a redox active disulfide bond. J Biol Chem 288, 34920–34929.

Higashi, S., Matsumoto, N., and Iwanaga, S. (1997). Conformation of factor VIIa stabilized by a labile disulfide bond (Cys-310-Cys-329) in the protease domain is essential for interaction with tissue factor. J Biol Chem 272, 25724–25730.

Hogg, P.J. (2013). Targeting allosteric disulphide bonds in cancer. Nature reviews Cancer 13, 425–431.

Hopfner, K.P., Lang, A., Karcher, A., Sichler, K., Kopetzki, E., Brandstetter, H., Huber, R., Bode, W., and Engh, R.A. (1999). Coagulation factor IXa: the relaxed conformation of Tyr99 blocks substrate binding. Structure 7, 989–996.

Kaelin, A.C. (1975). Sodium periodate modification of factor VIII procoagulant activity. Br J Haematol 31, 349–359.

Knudsen, T.T., Thorsen, S., Jensen, S.A., Dalhoff, K., Schmidt, L.E., Becker, U., and Bendtsen, F. (2005). Effect of intravenous N-acetylcysteine infusion on haemostatic parameters in healthy subjects. Gut 54, 515–521.

Lackman, R.L., Jamieson, A.M., Griffith, J.M., Geuze, H., and Cresswell, P. (2007). Innate immune recognition triggers secretion of lysosomal enzymes by macrophages. Traffic 8, 1179–1189.

Manning, F., Fagain, C.O., O’Kennedy, R., and Woodhams, B. (1995). Effects of chemical modifiers on recombinant factor VIII activity. Thromb Res 80, 247–254.

McMullen, B.A., Fujikawa, K., Davie, E.W., Hedner, U., and Ezban, M. (1995). Locations of disulfide bonds and free cysteines in the heavy and light chains of recombinant human factor VIII (antihemophilic factor A). Protein Sci 4, 740–746.

Miyata, T., Kawabata, S., Iwanaga, S., Takahashi, I., Alving, B., and Saito, H. (1989). Coagulation factor XII (Hageman factor) Washington D.C.: inactive factor XIIa results from Cys-571----Ser substitution. Proc Natl Acad Sci U S A 86, 8319–8322.

Pasquarello, C., Sanchez, J.C., Hochstrasser, D.F., and Corthals, G.L. (2004). N-t-butyliodoacetamide and iodoacetanilide: two new cysteine alkylating reagents for relative quantitation of proteins. Rapid Commun Mass Spectrom 18, 117–127.

Ramos, C.R., Abreu, P.A., Nascimento, A.L., and Ho, P.L. (2004). A high-copy T7 Escherichia coli expression vector for the production of recombinant proteins with a minimal N-terminal His-tagged fusion peptide. Braz J Med Biol Res 37, 1103–1109.

Read, S.A., O’Connor, K.S., Suppiah, V., Ahlenstiel, C.L.E., Obeid, S., Cook, K.M., Cunningham, A., Douglas, M.W., Hogg, P.J., Booth, D., et al. (2017). Zinc is a potent and specific inhibitor of IFN-lambda 3 signalling. Nature Communications 8, 15245

Savidge, G., Carlebjork, G., Thorell, B., Hessel, B., Holmgren, A., and Blomback, B. (1979). Reduction of factor VIII and other coagulation factors by the thioredoxin system. Thromb Res 16, 587–599.

Schwertassek, U., Balmer, Y., Gutscher, M., Weingarten, L., Preuss, M., Engelhard, J., Winkler, M., and Dick, T.P. (2007a). Selective redox regulation of cytokine receptor signaling by extracellular thioredoxin-1. EMBO J 26, 3086–3097.

Schwertassek, U., Weingarten, L., and Dick, T.P. (2007b). Identification of redox-active cell-surface proteins by mechanism-based kinetic trapping. Sci STKE 2007, p18.

Yousef, G.M., Elliott, M.B., Kopolovic, A.D., Serry, E., and Diamandis, E.P. (2004). Sequence and evolutionary analysis of the human trypsin subfamily of serine peptidases. Biochim Biophys Acta 1698, 77–86.

